# The RNA-binding protein PRRC2B preserves 5’ TOP mRNA during starvation to maintain ribosome biogenesis during nutrient recovery

**DOI:** 10.1101/2024.12.04.626744

**Authors:** Nadav Goldberg, Doron Bril, Miriam Eisenstein, Tsviya Olender, Alon Savidor, Shani Bialik, Shmuel Pietrokovski, Adi Kimchi

## Abstract

PRRC2B is an intrinsically disordered RNA-binding protein that is part of the cell’s translation machinery. Here we show that PRRC2B has two alternatively spliced mRNA transcripts producing major long and minor short isoforms. Mass spectrometry-based interaction studies indicated that both isoforms associate with the 40S ribosomal subunit and translation initiation factors. Importantly, the long isoform also interacted with additional RNA-binding proteins through its unique Arg/Gly-rich region. Among these is LARP1, a regulator of 5’ terminal oligopyrimidine (TOP) mRNAs under conditions of mTOR inhibition. We discovered that like LARP1, PRRC2B is necessary for preservation of 5’ TOP mRNA levels, particularly those encoding ribosomal proteins, during amino acid starvation. In its absence, the rapid *de novo* translation of ribosomal proteins that takes place upon nutrient recovery is impeded. Overall, our study elucidates a newly discovered function for PRRC2B as an RNA-binding protein that regulates ribosomal biogenesis upon metabolic shift, in addition to its established function in initiating translation of specific mRNA targets.

## Introduction

RNA binding proteins (RBPs) are important regulators of gene expression and homeostasis. These proteins affect and accompany every step of the mRNA life cycle, from transcription, splicing and nuclear export, to localization, translation, stability and degradation (*1*). Their functions are important in normal growth conditions, and following various stresses, such as nutrient starvation, oxidative stress, and DNA damage (*2, 3*). RBPs are also crucial for embryonic development, pluripotency, differentiation, and tumorigenesis (*4*). The RNA interactome is now recognized as consisting of thousands of proteins, thanks to large scale mass spectrometry-based proteomic screens of RNA interacting proteins. This, combined with techniques involving UV-crosslinking and RNA sequencing, has uncovered both the target mRNAs and the interacting interface of many RBPs, greatly advancing our understanding of RNA-protein regulatory processes (*5*).

Proline-rich coiled-coil 2 (PRRC2) proteins are a family of RBPs comprised of paralogues PRRC2A, B and C, all of which are large intrinsically disordered proteins. All three were identified in unbiased interactome screens for RBPs (*6–12*). PRRC2A has been shown to be an m(6)A reader that can stabilize its target mRNA, Olig2, through binding a methylated consensus motif, thereby ultimately regulating oligodendrocyte development (*13, 14*). This property is shared by the family, and PRRC2B similarly stabilizes *SOX2* mRNA to facilitate oligodendrocyte progenitor cell development and myelination (*15*). In cerebral endothelial cells, PRRC2B also binds m(6)A containing mRNAs and regulates expression of ECM components by mechanisms involving mRNA splicing and mRNA decay (*16*). In addition, all 3 PRRC2 proteins localize to stress granules, and PRRC2C was shown to be necessary for stress granule formation (*17*). Notably, recent data is consistent with a role in translation initiation, as the PRRC2 proteins associate with translation initiation factors and the pre-initiation complex (PIC) and fractionate with both 40S ribosome and 43-48S PICs (*18*). Deletion of PRRC2 proteins from HeLa cells leads to decreased global translation rates and a corresponding decrease in proliferation (*18*). The phenotype is more severe with double or triple deletions/depletion, suggesting either redundant functions within the family, or a combined effect of the loss of specific targets of each family member. Ribosome footprinting of the triple KD cells indicated that PRRC2 proteins direct the translation of specific mRNA targets with long 5’UTRs that harbor upstream reading frames (uORFs). Moreover, PRRC2C selectively binds ribosomes on mRNAs with uORFs, correlating with those mRNAs that show decreased translation efficiency upon PRRC2 knock-down. Furthermore, PRRC2 was shown to promote leaky scanning through these uORFs, thereby driving translation of the main ORF (*18*). The *Drosophila* PRRC2 homologue, Nocte, was similarly shown to direct translation of mRNA targets with uORFs, but this was shown to be promoted by re-initiation downstream to the uORFs (*19*). PRRC2B was shown to bind mostly to the 5’ UTR of its target mRNAs, within proximity of the translation start site, specifically recognizing GA and CU-rich motifs (*20*). mRNAs whose translation require PRRC2B include oncogenes and cell cycle regulators, specifically, cyclin D (*20*). While these studies have significantly advanced our understanding of PRRC2 function, the disparate functions elucidated so far imply multiple roles for these proteins, and additional mRNA targets and RNA-regulatory functions remain to be discovered.

In this study we focus on PRRC2B and shed new light on its role as an RNA-regulatory protein. We identified two splice isoforms, a long and short form, and assessed their respective functions. While both forms interact with translation initiation factors and the 40S ribosome, the long form has a unique repertoire of interacting proteins that confer additional functions. Specifically, PRRC2B_L_ interacts with LARP1 and PABP, and PRRC2B is involved in the preferential preservation of TOP mRNA levels, in particular those encoding ribosomal proteins, during prolonged amino acid starvation. The depletion of PRRC2B during starvation prevents the maintenance of these mRNAs, resulting in inhibition of *de novo* translation of ribosomal proteins upon refeeding that may slow down the recovery of cells from metabolic stress.

## Results

### PRRC2B has two splice isoforms, PRRC2B_L_ and PRRC2B_S_

Western blotting of various human cell lines for PRRC2B expression with an antibody that recognizes the C-terminus of the protein (amino acids 2102-2139) revealed two bands of different sizes: a slower migrating band that corresponds to the expected protein size of 250 kDa, and a faster migrating, shorter form at around 180 kDa (Fig. 1A). We observed these forms in all cells assayed, including fibroblasts and cancer cells of different origins, at different proportions, with the long form predominating. To investigate the origin of these forms, we performed PCR on cDNA generated from HEK293T cells using primers from either end of the predicted coding sequence of PRRC2B. Two PCR products were detected, the longer of which matched the predicted size of the cDNA encoding the reported PRRC2B protein of 250 kDa (Fig. 1B). The smaller band was sequenced to determine its identity, specifying a cDNA encoding a protein identical to the reported PRRC2B, but lacking nucleotides 2325-4407 (amino acids 775-1468). We noted multiple alternative splicing events within the PRRC2B gene upon analysis of NCBI RefSeq gene products on the UCSC genome browser (https://genome.ucsc.edu/cgi-bin/hgGateway, human assembly GRCh38/hg38). Specifically, the smaller form sequenced in HEK293T cells corresponds to a splice variant lacking a single long exon (Exon 16) (Fig. 1C). Importantly, levels of both protein bands were reduced upon transfection of HEK293T cells with a pool of siRNA targeting regions common to both the long and short isoforms within the CDS and 3’ UTR (siBOTH), but only the longer protein showed reduced expression upon transfection of siRNA targeting the CDS within exon 16 (siEx16) (Fig. 1D). This confirms the identity of the protein isoforms, which we will hereafter refer to as PRRC2B-long (PRRC2B_L_) and PRRC2B-short (PRRC2B_S_), respectively.

**Figure 1.**
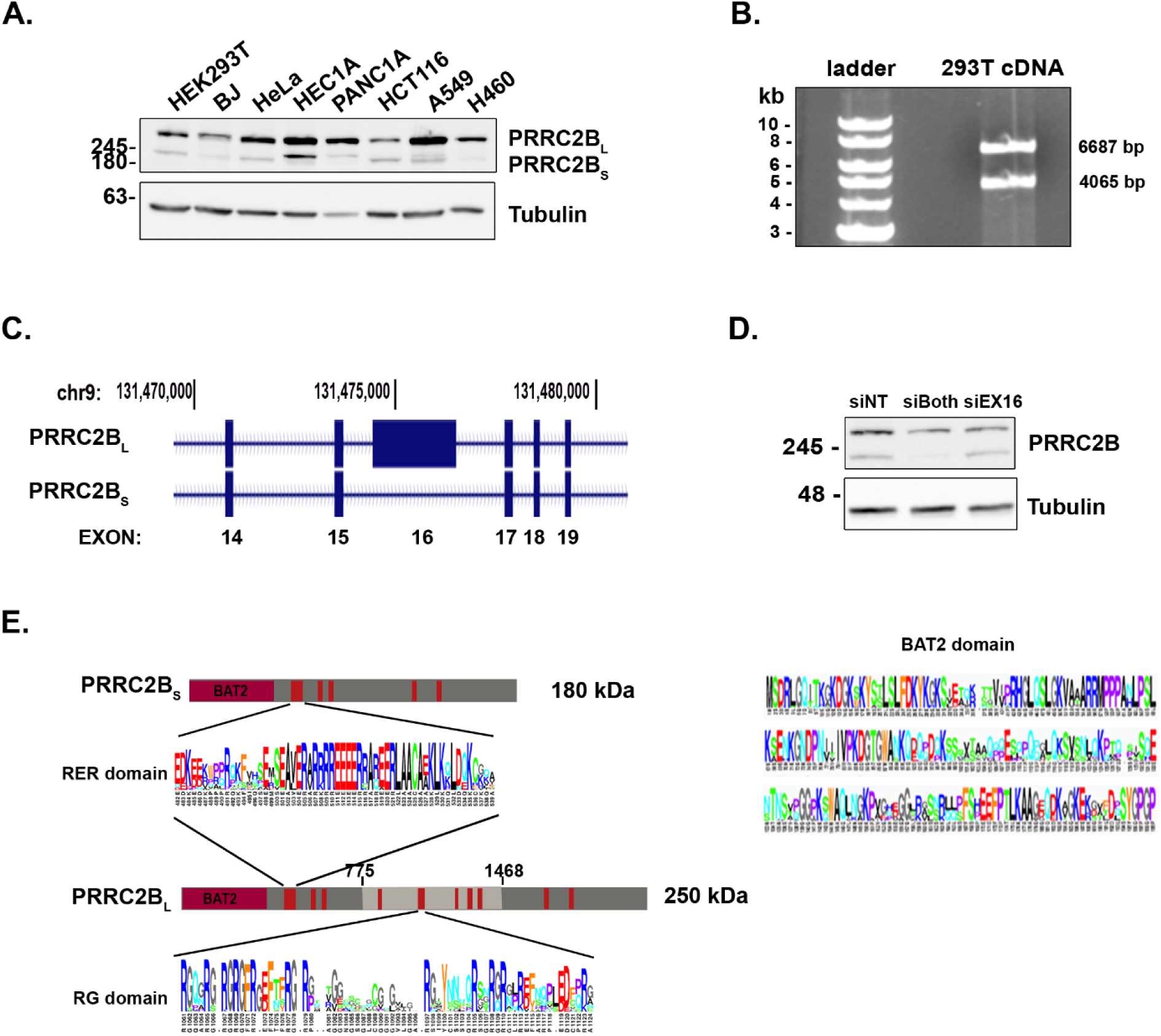
PRRC2B is expressed as long and short isoforms in human cell lines. (**A**) Western blots of the indicated cell lines for PRRC2B. Tubulin was used as a loading control. (**B**) Products of PCR performed on cDNA isolated from HEK 293T cells using primers from 5’ and 3’UTRs. (**C**) Portion of the PRRC2B genomic locus as represented on the UCSC genome browser depicting two splice isoforms, zooming in on exon 16 and surrounding exons. Numbers on top represent chromosomal position. (**D**) HEK 293T cells were transfected with control non-targeting (NT) siRNA, or siRNA to regions common to both isoforms (siBoth), or only exon 16 (siEx16), and after 3 days, lysates were subjected to western blot for PRRC2B and tubulin as loading control. (**E**) Schematics of PRRC2B_L_ and PRRC2B_S_ protein structures, with regions of high conservation colored red. Region derived from exon 16 in PRRC2B_L_ is colored light grey. Multiple sequence alignment within the indicated BAT2, RER and RG-rich domains are shown according to the human amino acid position numbers.

By this analysis, both PRRC2B_L_ and PRRC2B_S_ share the same N- and C-termini, the former consisting of a 200 amino acid conserved region termed the BAT2 domain (Fig. 1E) (*21*). However, the short isoform lacks the central part of the protein, including the conserved region encompassing aa 1061-1124, which contains 7 di-repeats of the amino acid Arg followed by Gly (referred to hereafter as the RG-rich domain). PRRC2B actually has 15 RG dipeptides throughout its structure, although this quantity is not statistically enriched compared to the expected random association of these residues, based on the total number of Arg and Gly residues present (as calculated based on (*22, 23*). However, the RG sites were significantly concentrated in exon 16 (right sided Fisher’s exact test (FET), *p*=0.001). Multiple alignments of this region across vertebrate families (Fig. 1E, bottom) showed that the RG-rich region is highly conserved (Mann-Whitney U test for whether the sites outside exon 16 are less conserved than those in it, after weight-correction for occurrence of RG sites in human proteins; *p*=0.037). Considering that conserved RGG/RG motifs are found in many RNA binding proteins, and have been shown to mediate interactions with both nucleotides and proteins (*24*), the significant enrichment and conservation of the RG-rich domain implies functional importance of this region.

### PRRC2B_L_ and PRRC2B_S_ share common structural and functional features

We used the Alphafold-3 (https://golgi.sandbox.google.com/, queries Q5JSZ5, Q5JSZ5-5) or RoseTTAfold servers (*25, 26*) to model the structures of both long and short PRRC2B isoforms (Fig. 2A, B). PRRC2B_L_ is predicted to be a mostly unstructured protein with almost no internal contacts or folded domains. The most prominent modeled feature is a long helix (residues 492-550) with a highly charged surface (Fig. 2A, bottom), bearing the conserved sequence RKRREEEERR (see Fig. 1E). This sequence (referred hereafter as the RER motif) is conserved in PRRC2A and PRRC2C as well (fig. S1A). Analysis of the sequence of PRRC2B by the prediction algorithm DisoRDPbind (http://biomine.cs.vcu.edu/servers/DisoRDPbind/) (*27*) showed several peaks with predicted RNA binding propensity, the strongest of which corresponded to this region (fig. S1B). This helix is surrounded by shorter helices and unstructured coils that differ in length and position in the 5 models predicted by Alphafold or RoseTTAfold. None of them contact the long helix, hence a folded domain is not formed. PRRC2B_s_ is similarly unstructured, with the long charged α-helix prominent in its center, although the BAT2 domain and the other smaller helices are differentially positioned compared to PRRC2B_L_ (Fig. 2B). These predictions are consistent with previous descriptions of PRRC2 family members as intrinsically disordered proteins (*19*), a feature common to RNA binding proteins and proteins that form membrane-less organelles.

**Figure 2.**
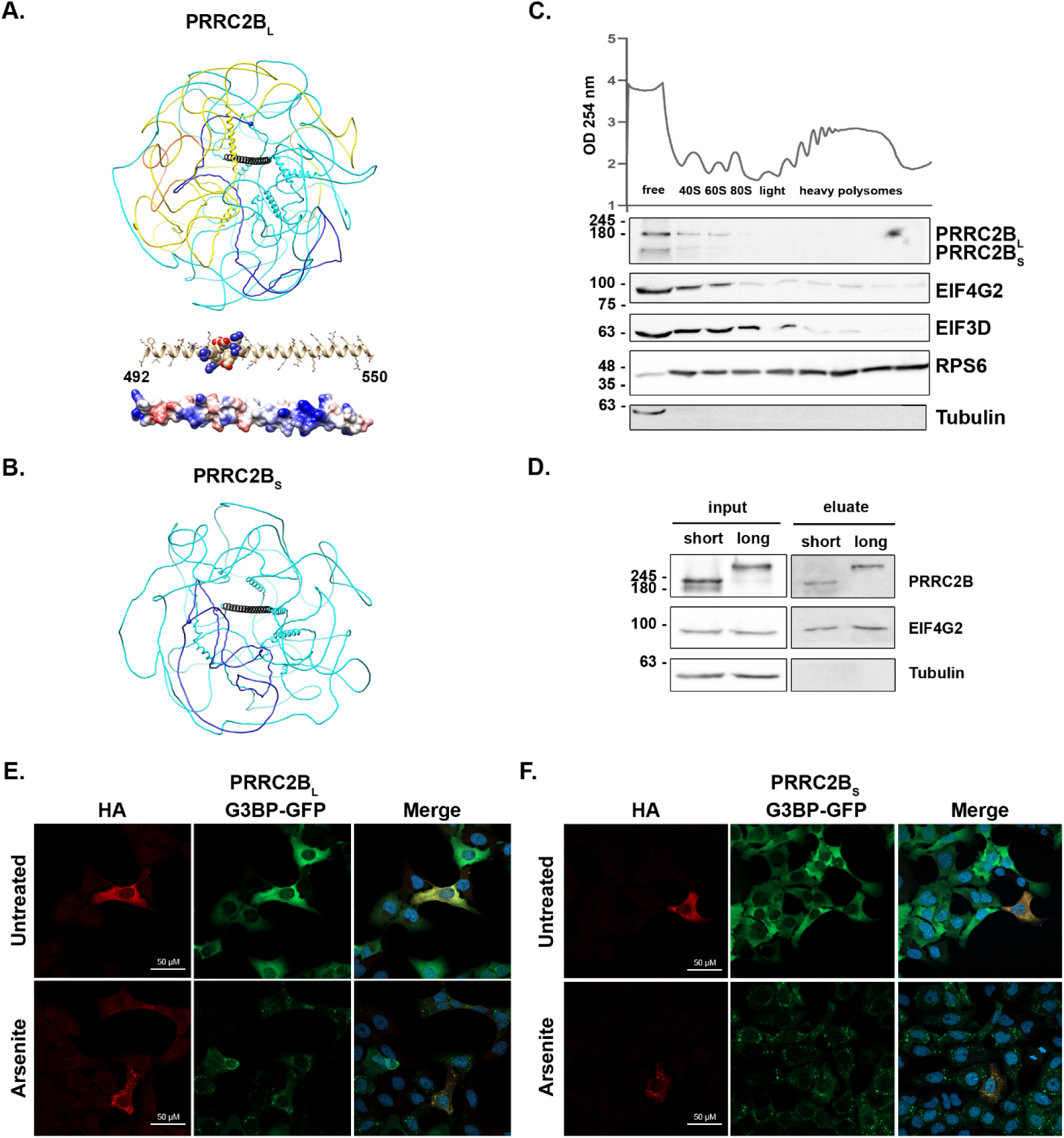
Both PRRC2B isoforms exhibit features of translation initiation factors. **(A, B)** Three-dimensional model structures of PRRC2B_L_ (A) and PRRC2B_S_ (B) according to predictions by Alphafold-3 server. Top AF-3 models are shown in cyan with the BAT2 domain (1–200) in blue. The N-terminus is indicated by the blue sphere. The region encoded by exon 16 that is lacking in the short isoform is colored yellow, with the RG-rich region 1061-1124 in red in the structure of PRRC2B_L_ shown in A. The long helix 492-550, shown in black in both structures, is also presented at the right in A as a ball-and-stick model with oxygen and nitrogen atoms in red and blue, respectively, and a solvent accessible surface colored by the Coulombic potential, with blue indicating positive, and red indicating negative surface regions, and white for neutral regions. Note that the highly conserved BAT2 segment is unstructured and does not form a folded domain. (**C**) Polysome profiling of HEK 293T cell lysate. Lysates were fractionated on sucrose gradients and 27 fractions were collected. Every 3 fractions were pooled and subjected to western blotting for the indicated proteins. (**D**) Oligo-dT pull down assay on HEK 293T lysates expressing HA-tagged PRRC2B_L_ or PRRC2B_S_. Lysates were cross-linked to stabilize RNA-protein complexes, and mRNAs with bound proteins were then captured on oligo-dT beads. Proteins present in the total cell lysate (input) and bead eluate were probed with anti-HA antibodies to detect PRRC2B. Endogenous EIF4G2 and Tubulin were used as positive and negative controls, respectively. (**E, F**) U2OS cells stably expressing GFP-G3BP were transfected with HA-tagged PRRC2B_L_ (E) or PRRC2B_S_ (F) and treated with sodium arsenite or DDW for 30 min. Cells were fixed and stained with anti-HA and DAPI to detect nuclei.

Previous reports have indicated that PRRC2B functions as a translation initiation factor (*18–20*). To explore whether the loss of exon 16 in PRRC2B_S_ confers functional differences to the isoform in this regard, we performed polysome purification and fractionation on lysates expressing both long and short PRRC2B. Notably, both isoforms were detected in association with free ribosomal subunits, particularly the 40S fractions, but not with polysomes, similar to other canonical and non-canonical translation initiation factors, such as EIF3D and EIF4G2, respectively (Fig. 2C). Furthermore, despite the absence of the RG-rich RNA binding domain in PRRC2B_S_, both isoforms were capable of binding mRNA, to similar extents as the non-canonical translation initiation factor EIF4G2, as determined from their presence among cross-linked mRNA-protein complexes that bound oligo-dT beads (Fig. 2D). Notably, mRNA binding persisted even when the RER motif was deleted from PRRC2B_S_ (fig. S1C), indicating that this motif is also not absolutely necessary for global interaction with mRNAs.

PRRC2 family members were previously shown to be components of stress granules (*17*), membraneless organelles acting as reservoirs of stalled translation initiation complexes, including 40S ribosome, translation initiation factors, mRNAs and RBPs, that accumulate during cell stress. To determine whether both isoforms of PRRC2B shift to stress granules in response to sodium arsenite treatment, we cloned long and short isoforms individually into pcDNA expression plasmids, fused to HA tag at the C-termini, and transfected them into U2OS cells expressing GFP-tagged G3BP, a stress granule component. Cells were then treated with sodium arsenite to induce stress. Staining for HA indicated that both PRRC2B isoforms shifted from a cytoplasmic diffuse localization under basal conditions to GFP-marked, punctate stress granules upon treatment of cells with sodium arsenite (Fig. 2E, F). Similar results were observed with a variant of PRRC2B_L_ in which the RG-rich domain was deleted (fig. S1D). Thus, loss of the central region containing the RG domain does not impair PRRC2B_S_’s ability to globally bind mRNA and initiation complexes, and to translocate to stress granules.

### The long and short PRRC2B isoforms have different protein interactomes

To further investigate the functional differences between the two isoforms, we examined the protein interactome of each isoform. We expressed HA-tagged PRRC2B_L_ and PRRC2B_S_ at low levels in HEK293T cells to simulate endogenous protein expression levels (fig. S2A). HA-PRRC2B isoforms and interacting proteins were immunoprecipitated with anti-HA agarose beads, eluted and analyzed by LC-mass spectrometry (IP-MS). A quantitative comparison was performed between identified proteins in the IP elutions of the two HA-PRRC2B isoforms and the HA-mCherry control, and between one another. In total, 170 specific candidate interacting proteins (≥2 unique peptides, FDR <0.05, and >2-fold enrichment over control HA-cherry IP) were identified for the long isoform, and a smaller set of 38 proteins interacted with the short isoform, all of which also interacted with the long isoform (Table S1, Fig. 3A-C). The larger number of interactors for PRRC2B_L_ is presumably due to the necessity of the large middle portion of the protein derived from exon 16 for these interactions. The top interactor of both isoforms was the non-canonical translation initiation factor EIF4G2, followed by DHX29, an RNA helicase involved in translation initiation, and members of translation initiation factors EIF1 and 3 (Table S1, Fig. 3A, B). Many additional initiation factors showed a significant yet lower abundance in the respective IPs (Table S1, Fig. 3A, B). Notably, both isoforms bound cytosolic ribosomal proteins, mostly of the small ribosomal subunit, consistent with their co-fractionation with 40S ribosomes observed above (Fig. 2C). Interestingly, even among the common interactors, the abundance of the initiation factors in the PRRC2B_L_ IP was roughly twice that of PRRC2B_S_ IP (Table S1). Examining those proteins that preferentially interacted with PRRC2B_L_ but not PRRC2B_S_ by focusing on those that were enriched in abundance more than 2-fold in the former IP (*p*<0.05), yielded a more focused set of 69 proteins. These mostly clustered into two main groups of proteins, as seen by network String analysis (Fig. 3D) (*28*). One group consisted of RBPs, including m6a readers, RNA helicases, proteins mediating stress granule formation, and most interestingly, RBPs involved in selective mRNA stabilization associated with translation regulation, such as the PABP family members PABPC1 and PABPC4, LARP1, LARP4 and LARP4B. The Arg methyltransferases, PRMT1 and PRMT5, were also identified in the list of proteins that preferentially bind to PRRC2B_L_. The second group consisted of 14 proteins of the mitochondrial ribosomal small subunit (MRPS). This is quite surprising, as PRRC2B_L_ localizes to the cytosol with no signs of intramitochondrial staining (fig. S2B), suggesting interactions with MRPS occur in a cytosolic compartment prior to their mitochondrial translocation. Pathway analysis of the set of 69 proteins that preferentially interact with PRRC2B_L_ by GeneAnalytics GO Biological Processes confirmed these functional classifications (fig. S2C).

**Figure 3.**
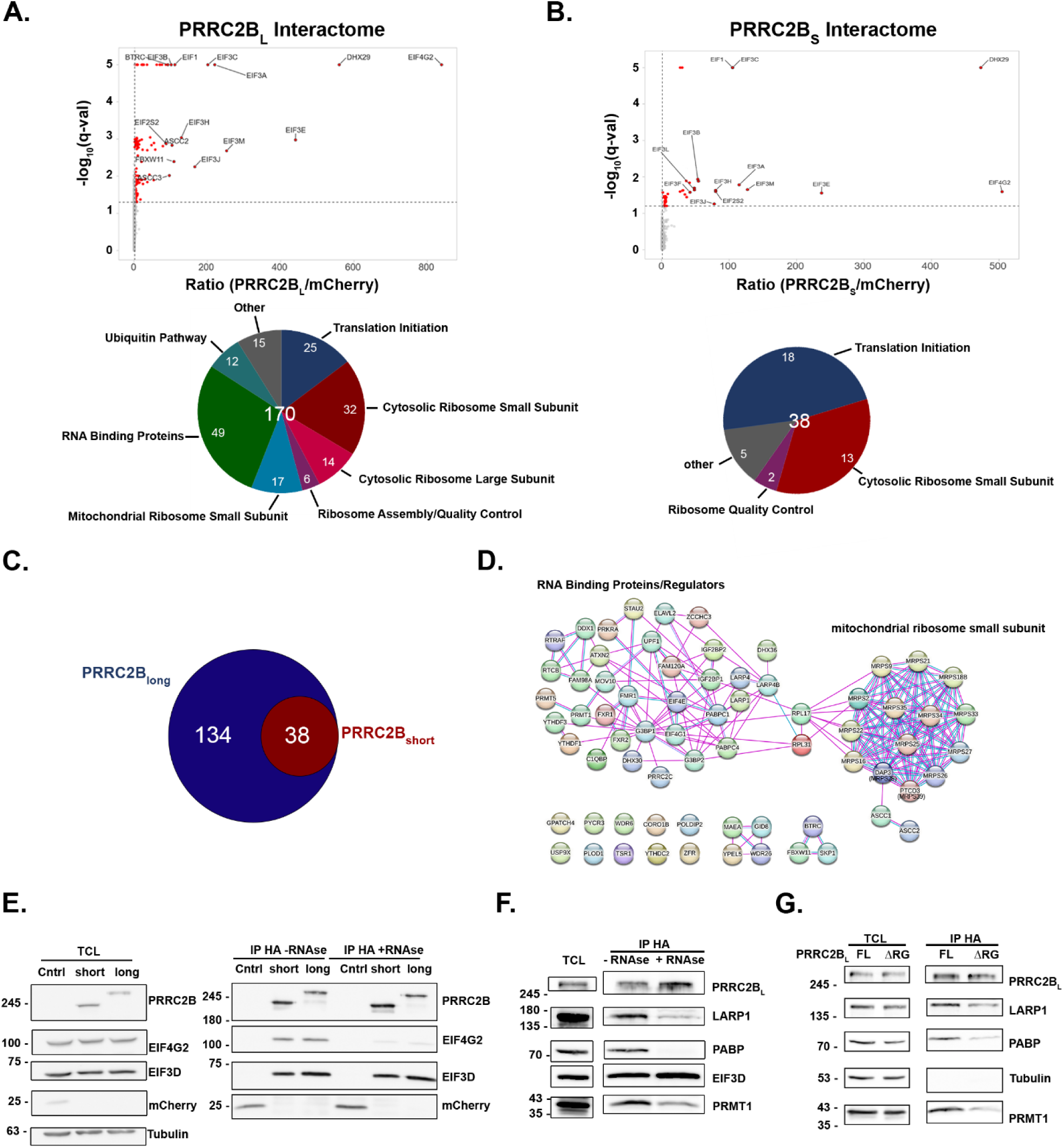
PRRC2B_L_ and PRRC2B_S_ have different protein interactomes. (**A-D**) HEK 293T cells were transiently transfected with pcDNA plasmids expressing HA-mCherry as control, HA-tagged PRRC2B_L_ or PRRC2B_S_. 24h after transfection HA-protein complexes were co-immunoprecipitated and analyzed by MS. (**A, B**) Volcano plot of the fold-ratio of the abundance of the detected proteins in PRRC2B_L_ (A) or PRRC2B_S_ (B) vs control IP samples, vs. their significance expressed as -log_10_ *q*-value, Student’s T-test. Top proteins with significant increased abundance are indicated in red for each isoform, highlighting translation initiation factors. Pie charts beneath graphs indicate distribution of interacting proteins within indicated categories. (**C**) Venn diagram depicting overlap between PRRC2B_L_ and PRRC2B_S_ interactomes. (**D**) String network analysis of set of 69 proteins that show significant enrichment (FC>2, *p*< 0.05) in PRRC2B_L_ interactome compared to PRRC2B_S_ interactome. (**E, F**) HEK 293T cells transiently transfected with pcDNA plasmids expressing HA-mCherry as control, HA-tagged PRRC2B_L_ or PRRC2B_S_ were immunoprecipitated with anti-HA antibodies in the absence of presence of 100μg/ml RNase A, and lysates and IPs subjected to western blot analysis for the indicated proteins. (**G**) HEK 293T cells transiently transfected with pcDNA plasmids expressing HA-tagged PRRC2B_L_ full length or ΔRG mutant were immunoprecipitated with anti-HA antibodies, and lysates and IPs subjected to western blot analysis for the indicated proteins.

We validated the interactions of EIF3D and EIF4G2, representative of translation initiation factors, with both long and short PRRC2B isoforms by co-IP/western blots. Both isoforms co-IPed with EIF4G2 and EIF3D (Fig. 3E), an interaction that required the common NH_2_-terminal BAT2 domain (fig. S2D). Interestingly, immunoprecipitation in the presence of RNase A reduced the interaction with EIF4G2, but not with EIF3D (Fig. 3E), indicating that the former requires RNA. In addition, several of the exclusive PRRC2B_L_ interacting proteins were also validated by co-IP/western blot, including LARP1, PABP and PRMT1 (Fig. 3F). All three proteins co-immunoprecipitated with PRRC2B_L_, and in contrast with EIF3D, the interactions were decreased upon RNase A treatment. Notably, and in agreement with the MS data, the co-IP between PRRC2B_S_ and LARP1 was greatly reduced compared to that of PRRC2B_L_ (fig. S2E). Deletion of the RG-rich domain, the putative RNA binding region, from PRRC2B_L_ also reduced its binding to LARP1, PABP and PRMT1 (Fig. 3G). The latter is particularly interesting as R1077 and R1079 within PRRC2B_L_’s RG-rich domain were identified as methylated by MS (fig. S2F, Table S2). As these are predicted substrates for methylation by PRMT1, the main mammalian type I Arg methyltransferase (*29*), it is notable that this region is both required for interaction with PRMT1 and likely modified by the methylase.

Altogether, the differences in interactomes suggest that the short PRRC2B isoform has more limited functional capability compared to the long isoform, resulting from expression of exon 16 in the latter.

### PRRC2B knock-out reduces TOP mRNA levels during amino acid starvation

Among the PRRC2B_L_ interactors were LARP1, PABPC1 and PABPC4, RBPs that are involved in the regulation of TOP mRNA stability and translation. TOP mRNAs, which include most cytosolic ribosomal proteins and several translation factors, bear a characteristic 5’ terminal oligopyrimidine (TOP) motif. This motif enables especial sensitivity to mTORC1 regulation, enabling efficient and immediate repression of translation initiation upon mTOR inhibition or inactivation (*30, 31*). Under basal conditions, when mTOR is active, TOP mRNAs are efficiently translated, and LARP1 binds to the 3’UTR of these mRNAs together with PABP. Upon the inhibition or inactivation of mTOR, such as following amino acid starvation, LARP1 is dephosphorylated and interacts also with the TOP motif and the m(7)G 5’cap, thereby blocking EIF4E cap binding, and consequently shutting down cap-dependent translation (*31–36*). At the same time, LARP1 binding to the polyA tail (*37*) specifically preserves TOP mRNA levels following mTOR inhibition as a complex with the 40S ribosome, by promoting TOP mRNA stability (*38–40*). Moreover, it facilitates post-transcription polyA elongation during amino acid starvation, and as a result preserves pools of TOP mRNA for immediate ribosome loading when metabolism is restored (*41*). Upon re-feeding and recovery of mTORC1 activity, LARP1 is immediately phosphorylated by mTOR, and its release from the 5’ UTR of TOP mRNAs enables their translation by cap-dependent factors.

Interestingly, PRRC2B was shown to specifically bind CU rich sequences within the 5’ UTR of its interacting mRNAs (*20*), similar to LARP1 (*42*). Considering the specific interactions between PRRC2B_L_ and LARP1 and PABP, we hypothesized that PRRC2B_L_ may be an additional RBP that participates in LARP1-mediated mRNA regulation. Moreover, GO annotation of the list of PRRC2B_L_ interacting proteins for enriched biological processes included the term “regulation of mRNA stability” (fig S2C). To assess PRRC2B’s potential role in regulating TOP mRNA stability, we used CRISPR-Cas9 to knock-out (KO) both isoforms of PRRC2B in HEK293T cells, using a non-targeting guide to generate control cells (NT) (Fig. 4A).

**Figure 4.**
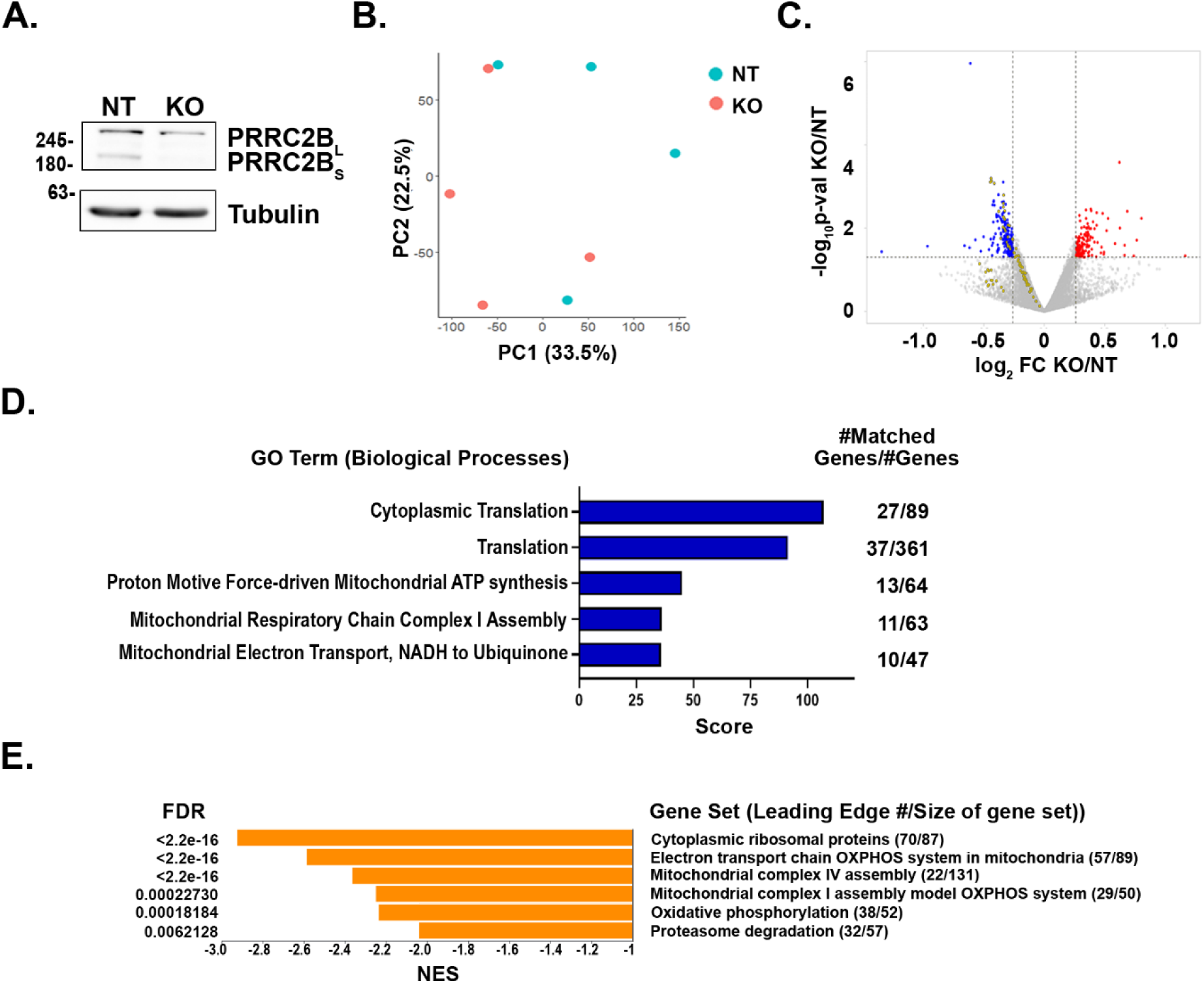
PRRC2B is necessary for maintenance of TOP mRNA expression during starvation. (**A**) HEK293T PRRC2B KO and control NT KO cells were generated by Cas9-CRISPR. Western blotting of pooled polyclonal KOs confirm reduction in PRRC2B expression. Tubulin was used as a loading control. (**B-E**) RNA-seq was performed on PRRC2B KO and NT cells in basal conditions and following 48h starvation in EBSS media. (**B**) Principle Component Analysis (PCA) of RNA seq data from starved NT or PRRC2B KO cells. (**C**) Volcano plot of the fold-change in mRNA expression (protein-coding only) levels in PRRC2B KO vs NT control under starvation conditions, vs. their significance expressed as -log_10_ *p*-value. Increased and decreased differentially expressed genes are indicated in red and blue, respectively. Yellow dots represent mRNAs encoding ribosomal proteins. (**D**) Venn diagram showing overlap between TOP mRNAs encoding ribosomal proteins (RPs) and the group of statistically significant down-regulated genes (down DEGs). (**E**) The set of 211 significantly decreased differentially expressed genes in starved PRRC2B KO cells vs control NT cells were analyzed by GeneAnalytics. Top 5 significant GO terms (Biological Processes) and scores are presented. The number of matched genes within the set out of the total number of genes in the particular biological process is indicated at right of graph. (**F**) The entire RNA-seq dataset was analyzed by WEB-based GEne SeT AnaLysis Toolkit (WebGestalt) gene set enrichment analysis (GSEA, https://www.webgestalt.org). Shown are the significant gene sets identified within the dataset, all of which have negative normalized enrichment scores (NES). The number of genes within the dataset (leading edge) out of the total number of genes in the particular gene set is indicated at right of graph.

To determine whether PRRC2B is necessary to maintain TOP mRNA levels during prolonged starvation, as demonstrated for LARP1, we performed RNA-seq on NT and PRRC2B KO HEK293T cells in basal conditions and following 48h amino acid starvation in EBSS media, focusing the analysis specifically on ribosomal and other TOP mRNAs (Table S3). Principle component analysis (PCA) indicated reproducibility of the replicates, which segregated mainly according to growth conditions (fig. S3A). First we confirmed, as previously reported (*38–40*), that despite the global decrease in mRNA levels following starvation, levels of TOP mRNAs were maintained relative to the majority of mRNAs (fig. S3B, Table S3). Technically, the calculations were based on loading of equal RNA amounts for the RNA-seq protocol and normalization of reads during the analysis. Thus, the genes whose levels decreased in EBSS samples were artificially scored as unchanged versus their levels in basal conditions. As a result, those genes whose mRNA expression dropped to a smaller extent than average, or did not actually change, were defined by the analysis as **increased** DEGs, along with those that actually increased. Notably, 90 out of 111 (81%, *p*<0.0001 by FET contingency analysis) mRNAs present in the RNA-seq dataset that were previously classified as TOP mRNAs (*30, 31*) were “increased” by more than 20% (*p*<0.05) (fig. S3B, Table S3), confirming that TOP mRNA expression is maintained during amino acid starvation in our cells. Focusing only on the genes encoding ribosomal proteins (RPs), 75 out of 77 (97%) that were detected in the dataset exhibited increased mRNA expression (*p*<0.0001 by FET contingency analysis). This suggests that in our system, RPs are more tightly regulated by starvation than other TOP genes.

To determine whether PRRC2B contributes to this regulation, we compared mRNA levels in NT and PRRC2B KO starved (EBSS) cells. Globally, there was a small degree of change in mRNA levels, as supported by the PCA analysis of the dataset, which showed no differences between the clustering of KO and NT replicates (Fig. 4B). A group of 211 genes, excluding PRRC2B, showed a significant (*p*<0.05) decreased expression of at least 20% (Fig. 4C, Table S3). Specifically, an examination of the 77 genes encoding ribosomal proteins that were detected in the dataset indicated that all were reduced to some degree, 35% (27/77) significantly (Fig. 4C). Venn analysis indicated that this represents a significant enrichment within the set of mRNAs with decreased expression (Fig. 4D). The effect was somewhat smaller when examining all TOP mRNAs; out of the 111 reported TOP mRNAs detected, 31 (28%) showed significantly reduced abundance (fig. S3C). Again, this was a significant enrichment (Fig. 4D). GeneAnalytics GO analysis of the group of significantly down-regulated mRNAs yielded 18 categories with similar gene components that were reiterations on either translation or mitochondrial oxidative respiration/ATP generation (top 5 are shown in Fig. 4E). GSEA analysis of the dataset, which assesses even small changes that become significant when occurring within multiple components of the same pathway, also highlighted the ribosomal proteins as significantly downregulated (Fig. 4F). Thus, the RNA-seq analysis indicates that the mechanism that maintains the expression of TOP mRNAs during starvation is impaired upon KO of PRRC2B, particularly for those encoding ribosomal proteins.

### PRRC2B is required for immediate translation of ribosomal proteins during post-starvation recovery

The persistence of mRNAs encoding ribosomal proteins during prolonged amino acid starvation enables their immediate translation following re-feeding as they are prepared for ribosomal loading (*41*). Thus, by impairing this persistence PRRC2B KO should also affect the first wave of translation of these mRNAs upon nutrient restoration. To assess this, we measured PRRC2B KO’s effects on *de novo* synthesis of ribosomal proteins during basal, amino acid starvation (EBSS) and nutrient restoration (re-feeding) conditions. Nascent proteins of NT and PRRC2B KO HEK293T cells were labeled with AHA, a Met analogue that can be isolated and detected via a biotin moiety that is subsequently attached by CLICK-IT chemistry (Fig. 5A). Western blotting of labeled proteins for RPS3, as a representative TOP mRNA-encoded protein, and non-TOP protein calnexin, showed reduction in both proteins following 48h of amino acid starvation. The latter reflects global mechanisms that reduce cap-dependent translation in response to mTOR inhibition (e.g., 4EBP dephosphorylation) (*3*). The reduction of RPS3 translation was consistently stronger, due to the additional suppressive mechanism imposed by LARP1 dephosphorylation described above (*31–36*) (Fig. 5B). PRRC2B KO did not affect translation repression during starvation. Resynthesis of both proteins was observed 1h after restoration of nutrients, with the ribosomal protein showing the most robust recovery, consistent with its prioritized translation (*39*). Most importantly, the *de novo* synthesis of RPS3 protein following re-feeding was reduced in PRRC2B KO cells (Fig. 5B, C). The effect of PRRC2B KO on non-TOP protein translation is milder at this early time. Thus, PRRC2B is required for the immediate translation of TOP mRNAs such as RPS3 following restoration of nutrients.

**Figure 5.**
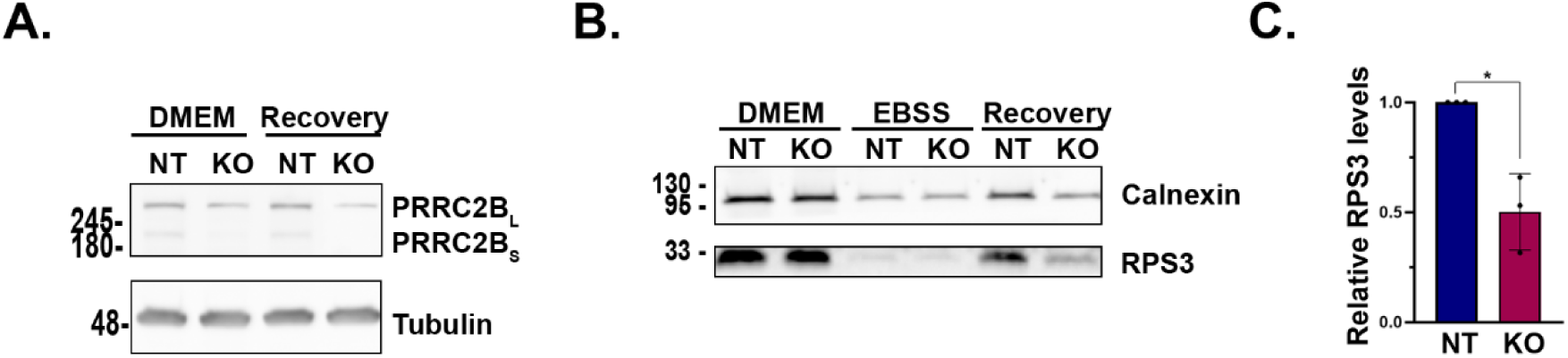
PRRC2B is necessary for resumption of translation of ribosomal proteins during recovery from starvation. (**A**) Western blot showing levels of PRRC2B and tubulin, as loading control, in basal conditions and 1h of restoration of media following 48h starvation (Recovery). (**B, C**) PRRC2B KO and NT control cells under normal growth conditions, following 48h starvation (EBSS) and after 1h restoration of met free growth media (Recovery) were incubated with Met analogue AHA, which was then attached to biotin by Click chemistry. Biotin-labeled nascent proteins were purified from total cell lysates on streptavidin beads and subjected to western blot with the indicated antibodies (B). Shown is a representative blot of n=3 experiments. (C) RPS3 levels during recovery were normalized to the non-TOP protein calnexin. Data is presented as means±SD of 3 experiments, statistical significance was determined by 2-tailed T-test, with unequal variance. *, *p* = 0.038.

## Discussion

Here we identify two expressed isoforms of the RNA binding protein PRRC2B, a long form, PRRC2B_L_, and a shorter form, PRRC2B_S_, lacking the middle region of the protein. Our data indicate that both PRRC2B_L_ and PRRC2B_S_ bind multiple translation initiation factors, interact with and co-fractionate with the 40S ribosome, and bind mRNAs. The common N- and C-termini that are shared by both proteins have been shown to be important for protein binding; the N-terminus in particular has been shown to bind translation initiation factors (*19, 20*). We confirm the role of the N-terminal BAT2 domain in interacting with both canonical and non-canonical translation initiation factors, and show that this characteristic is common to both PRRC2B isoforms. By assessing the effect of RNase treatment on the presence of proteins in these co IPs, we distinguished between direct and indirect protein-protein interactions. For example, while EIF3D interacts with PRRC2B even in the presence of RNase, EIF4G2’s interaction with PRRC2B was strongly suppressed, suggesting that the presence of RNA enables or stabilizes the interaction. Interestingly, both long and short isoforms are diverted to stress granules following arsenic stress. As only PRRC2B_L_ interacts with stress granule assembly proteins such FMR1, G3BP1, and G3BP2, the translocation of PRRC2Bs presumably results from its association with stalled translation initiation complexes that accumulate in the stress granules. Based on the similar behavior demonstrated for PRRC2B_L_ and PRRC2B_S_ in binding translation initiation factors, 40S ribosome and mRNA, we predict that both isoforms function as translation initiation regulators.

PRRC2B_S_ was present in the eluate bound to oligo-dT beads, indicating that it too can interact with mRNA, despite the fact that it lacks the middle of the protein that was shown previously to bind RNA (residues 750-1500) (*20*). This region was likewise shown to be involved in mRNA binding and translation activity in the *Drosophila* paralog Nocte (*19*). It is presumably the RG-rich region of this domain that mediates RNA binding. It has previously been proposed that the Gly residues permit flexibility of the protein backbone, which enables conformational plasticity of the disordered region. The positively charged Arg residues, on the other hand, provide non-specific electrostatic interactions with the mRNA and also more specific hydrogen bonding with mRNA tertiary structures (*43*). Thus overall, the RG-dipeptide motifs enhance the interaction with mRNAs. The RER motif following the N-terminal BAT2 domain may also potentially mediate RNA interactions, yet its deletion from PRRC2B_S_ did not prevent its interaction with mRNA. The persistence of mRNA binding by PRRC2B_S_ even in the absence of the middle RNA binding domain and the RER motif indicates that additional regions of the protein mediate the interaction. This is consistent with RNA-binding properties of intrinsically disordered RBPs that lack the classical RNA-binding domains, such as K-homology domains, Zinc fingers and RNA Recognition Motifs (RRM), and instead contain multiple RNA interaction sites of low complexity (*5, 44*). The multiple sites combined with the lack of an established, stable structure may enable greater adaptability and specificity in substrate recognition. Thus, the RG-rich and RER domains, although not necessary for global mRNA binding, may be critical for enhancing interactions with specific mRNA targets yet to be identified.

The functional differences between the two isoforms can be discerned from their different interactomes, as determined by the IP-MS proteomics. There are two major groups of proteins that preferentially interact with PRRC2B_L_ compared to PRRC2B_S_: the mitochondrial small ribosome subunit and RNA binding regulatory proteins. These interactions require the middle domain of the protein unique to PRRC2B_L_. We have shown that at least some of the proteins within the second group such as LARP1 and PABP, require the presence of the RG-rich region for efficient interaction. Moreover, treatment with RNase reduced the interaction, implying that these protein-protein interactions depend on PRRC2B’s interaction with RNA. It is possible that the interaction is indirect, i.e., both proteins bind the same mRNA simultaneously, which brings the 2 proteins into the same RNA-protein complex from which they co-IP. Alternatively, binding to mRNA may induce refolding of the intrinsically disordered PRRC2B, as such disordered proteins can become structured upon binding to other proteins or DNA/RNA (*45*), and in its newly acquired structured form the protein gains the ability to bind its interacting partner. Each of these two models can accommodate the possibility of functional interactions between these RNA binding proteins in a given RNA-protein complex.

The interaction with multiple protein components of the small subunit of the mitochondrial ribosome is intriguing, as PRRC2B does not localize to the mitochondria matrix where mitochondrial translation occurs. Furthermore, and in contrast to translation initiation factors of the cytosol that strongly interacted with PRRC2B, mitochondrial initiation factors were not part of the PRRC2B interactome. Therefore, PRRC2B is unlikely to play a parallel role in translation initiation of mitochondrial-encoded mRNAs within the mitochondria. Presumably, the interaction occurs in the cytosol. The interaction between PRRC2B_L_ and MRPSs may reflect a potential interaction that occurs when the latter accumulate in the cytosol, such as during import failure or mitochondria stress and rupture. As a protein that regulates translation initiation, PRRC2B may function as part of a quality control mechanism to sense the accumulating MRPSs and signal to the translation machinery to halt further translation of these proteins, until mitochondrial homeostasis is restored. A second possible explanation for the interaction may be derived from our finding on PRRC2B_L_’s interaction with the RNA binding LARP4. LARP4, like LARP1, selectively binds to specific cohorts of mRNAs to regulate their translation (*46*). It specifically binds to mRNAs encoding MRPs and proteins of the mitochondrial oxidative respiratory, in proximity of mitochondria (*47*). LARP4 is recruited to this cellular compartment by interacting with dAKAP1, a PKA adaptor scaffold protein that localizes to the mitochondrial outer membrane (*48*). Notably, dAKAP1’s tudor domain was also previously shown to interact with PRRC2 proteins (*48*). It is tempting therefore to speculate that PRRC2B is also part of this complex, interacting with the MRPSs as they are locally translated in proximity of the mitochondria prior to their import. Our current study does not address the mechanistic and functional significance of PRRC2B interactions with MRPSs, and additional investigation is required to determine if either of these hypotheses are correct.

PRRC2B_L_’s specific interaction with LARP1 and PABP proteins suggests a novel function for the protein in regulating mRNA stability of TOP mRNAs. We have shown that PRRC2B is necessary for maintaining levels of TOP mRNAs in general, and particularly ribosomal protein mRNA, during prolonged amino acid starvation conditions. PRRC2B’s necessity for maintaining RP mRNAs during starvation is evident in the impaired recovery of nascent ribosomal protein synthesis following restoration of nutrients. Re-synthesis of ribosomal proteins within the first hour of nutrient restoration is compromised in PRRC2B KO cells. We expect that this would likewise delay the recovery of ribosomal biogenesis, and as a consequence, the increase in global protein synthesis that occurs during these metabolic shifts. Thus, in addition to its previously demonstrated role in translation initiation under basal conditions (*18, 20*), PRRC2B likely indirectly affects translation by regulating levels of ribosomal protein biogenesis following stress.

PRRC2B is thus a new RBP player participating in TOP mRNA regulation during conditions of mTOR inactivation, along with LARP1 and PABP. Like the interaction between LARP1 and PABP (*42, 49*), the interaction between PRRC2B and LARP1 or PABP requires the presence of mRNA. Thus, one can envision all three proteins occupying positions on TOP mRNAs, which brings them into one ribonucleoprotein complex. One proposed mechanism by which TOP mRNAs are maintained during starvation is that they are preserved from degradation as part of a complex with the 40S ribosome (*39, 40*). It is tempting to speculate that PRRC2B, which can interact with ribosomal proteins, is responsible for recruitment of the 40S ribosome to the TOP mRNAs for this purpose. Yet another study showed that TOP mRNA preservation during starvation involves another role of LARP1 in shifting TOP mRNAs from polysomes to monosomes, from which low levels of translation occur to preserve ribosome thresholds (*50*). Although we have not specifically addressed this mechanism, PRRC2B would be well positioned for recruitment of the 80S ribosome to TOP mRNAs to initiate these low levels of translation, especially considering that the canonical EIF4G1- and cap-dependent initiation is inhibited upon mTOR inactivation. In either scenario, upon nutrient restoration, PRRC2B may continue its function in ribosome recruitment to TOP mRNAs, ensuring rapid recovery of proteins critical for further translation even prior to re-phosphorylation of EIF4EBP by mTOR and release of EIF4E for binding to the 5’ mRNA cap. It is notable that the LARP1-PRRC2B interaction was initially observed under basal conditions, even though PRRC2B KD did not affect *de novo* RP synthesis in basal media. Thus, while it may be capable of binding TOP mRNAs together with LARP1 and recruiting the ribosome, its contribution to initiation mechanisms is presumably less impactful, as cap-dependent initiation, involving functional EIF4E and EIF4G1, predominates in these circumstances. While the details of this working model await clarification by future research, the presence of an additional RBP that regulates TOP mRNA shed light on previously described LARP1-dependent mechanisms.

## Materials and Methods

### Cell lines and transfections

HEK293T cells (ATCC, cat# CRL-3216) were maintained in Dulbecco’s Modified Eagle’s Medium (DMEM; Sartorius cat# 01-055-1A) supplemented with 2mM glutamine (Gibco BRL), 100U/ml penicillin and streptomycin (Gibco BRL) and 10% fetal bovine serum (Gibco BRL). For starvation experiments, cells were washed in PBS and grown for 48h in EBSS media (Sartorius cat# 02-010-1A). U2OS cells expressing GFP-G3BP (*51*) were obtained from Prof. Eran Hornstein (Weizmann Institute) and grown in (DMEM). Additional cell lines used for western blot expression analysis were grown in DMEM. Cell lines were routinely tested for mycoplasma contamination and when relevant, authenticated by STR profiling.

Plasmids for expression of exogenous proteins were introduced to cells by standard calcium-phosphate transfection method. Transient PRRC2B KD was generated by transfecting HEK293T cells with control siRNA (Dharmacon D-001810-10) or siRNA targeting PRRC2B (Dharmacon L-032577-01-0005) using Lipofectamine 2000 transfection reagent (Invitrogen) according to manufacturer’s protocol. Sequences targeting both isoforms: TGAATGACCAAGACGGAAA (CDS), AGTGTAAGCAGGCACGAAA (CDS), CCACACAGCTCATCGTGAA 3’ (UTR). Sequence of siRNA targeting PRRC2B_L_ only (exon 16): 5’-GCTGAGCAATTGCGGGTAT-3’.

### Generation of PRRC2B KO Cells

PRRC2B KO HEK293T cells were generated using lentiviral mediated CRISPR-CAS9 as previously described (*52*). Briefly, guides targeting a region in exon 2 that contains the start codon (Guide 1: 5’-TTTCAAAGGCAGATCGGGAG-3’, Guide 2: 5’-GTAGACGCGATTAGATCCTC-3’) were cloned into lentcrispr V2 vectors carrying CAS9 and puromycin resistance genes. The guides target both isoforms as the N terminus is shared. A non-targeting guide (5’-TTTCGTGCCGATGTAACA-3’) was used for control KO. Virions were generated by standard methods. HEK293T cells were infected, following by selection with 1.5μg/ml Puromycin for 6 days. Genotyping was performed after selection using the following primers: Forward Primer 1: 5’-GTACTCCACAGATCGCCTCG-3’, Reverse Primer 2: 5’-TACATGCACACCCAGAAGCC-3’. Pools of infected cells were used for experiments; note that KO was not complete and depended on CRISPR efficiency within the pooled cells.

### Sequence alignment and motif analysis

PRRC2 protein sequences were identified in public sequence databases by blast searches starting with the human PRRC2B protein and then other identified vertebrate sequences. Sequences were multiply aligned using the GLAM2 (*53*) and blast programs, and multiple alignments were aligned to each other using the COMPASS program (*54*). Multiple alignment sequence logos visualization is as previously described (*55*). To quantify the occurrence of RG dipeptides in PRRC2B_L_ the product of the 164 occurrences of Arg and 200 occurrences of Gly was divided by the number of dipeptides in the PRRC2B_L_ protein (2228), giving a value of 14.72. This expected value is not significantly different from the observed value of 15 (*22, 23*). To test for the overabundance of RG dipeptides in exon 16, a right sided Fisher’s exact test (FET) was made with the number of RG dipeptides in exon 16 (11), the number not in exon 16 (4), the number of dipeptides in exon 16 (694), and the number not in exon 16 (1533).

### Cloning of PRRC2B constructs

RNA from HEK293T cells was purified using Monarch total RNA miniprep kit (NEB). cDNA was reverse-transcribed with SuperScript III (Invitrogen). Full length PRRC2B long and short isoforms were amplified from cDNA by PCR (KAPA HiFi HotStart Ready Mix PCR Kit) with primers containing restriction sites for KpnI and EcorV. The reverse primer also contained the HA tag sequence and two stop codons. The amplified products were then inserted into pcDNA expression vector using the forementioned restriction enzymes. PRRC2B ΔBAT2 was constructed using whole plasmid PCR with phosphorylated primers: Forward primer: 5’-ATGCTCCGCCCTCAGAATGTG-3’ Reverse primer: 5’-GGTACCAAGCTTGGGTCTCC-3’.

pcDNA PRRC2B_L_/_S_ plasmids were used as templates. The reaction was then treated by DpnI for 2 h to remove the original plasmid. The PCR product was cleaned using QIAquick PCR Purification Kit (Qiagen) and ligated into pCDNA3. ΔRG PRRC2B construct was created in the same manner using the following primers: Forward: 5-’ CTGCGAGAGTTTGCGCGGC-3’, Reverse: 5’-GACCCCAAAGGCTTGCTCC-3’.

### Co-immunoprecipitation

HEK293T cells were transiently transfected for 24 h with HA-tagged PRRC2B constructs or HA-mCherry as control. Cells were lysed in buffer B (20 mM HEPES-KOH [pH 7.6], 100 mM KCl, 0.5 mM EDTA, 0.4% NP-40, 20% glycerol) supplemented with protease and phosphatase inhibitors (Sigma), 0.1 mM phenylmethylsulfonyl fluoride (PMSF). 100μg/ml RNase A (Thermo Scientific, EN0531) was added to relevant samples. Lysates were incubated with anti-HA beads (Sigma) for 2 h at 4°C. Following 3 washes with buffer B, the beads were incubated for 5 min with Elution buffer (100mM Tris-HCl, 5% SDS PH 7.4) and analyzed by mass spectrometry or western blotting on 8% Tris-Glycine gels.

### Oligo-dT beads pulldown

Assays for mRNA binding were done according to previously described methods (*6*). In brief, HEK293T cells expressing PRRC2B isoforms were UV cross-linked (Stratagene Stratalinker UV 1800 Crosslinker) and 2 mg lysates were incubated with magnetic oligo-DT beads (NEB, S1419S) for 1 h. After stringent washes to remove non-specific and indirectly interacting proteins, bound mRNA-protein complexes were eluted from the beads and analyzed by western blotting.

### Click-iT labeling

*De novo* synthesized proteins were assayed as previously described (*41*). HEK293T PRRC2B KO and NT cells were grown in DMEM for basal conditions, or incubated in EBSS for 48h (starved), followed by 1h recovery in DMEM Methionine free media (Gibco, cat# 21013024) (recovery). For all conditions, cells were incubated in the appropriate methionine-free media (DMEM or EBSS) supplemented with 25 μM Click-iT AHA (Thermo Fisher Scientific, cat# C10102) for 1 h. Lysates were quantified using Pierce™ BCA Protein Assay (Thermo Scientific, cat# 23227) and between 80-100µg of proteins were labeled with biotin using the Click-iT protein reaction kit (Invitrogen, cat# C10276). Biotinylated proteins were precipitated according to manufacturer’s instructions and kept in −20° until further use. Proteins were resuspended, quantified again using BCA and equal amounts were either taken for total biotin signal or loaded on Pierce NeutrAvidin Agarose beads (Thermo Scientific™, cat# 29200) for 2 h at 4°C. Total biotin signal or immunoprecipitated biotinylated proteins were analyzed by western blotting for specific proteins as indicated in figure, or for total biotinylated proteins.

### Western Blot

Cells were lysed with B buffer or RIPA (20 mM Tris, pH 8.5, 0.1% NP40, 150mM NaCl, 0.5% sodium deoxycholate, 0.1% SDS) supplemented with 10 µl/ml 0.1M PMSF (Sigma-Aldrich, cat# 93482) and 1% protease inhibitor (Sigma-Aldrich, cat# P8340), and electrophoresis and western blots were performed according to standard protocols. Blots were stained with Ponceau S (Sigma, cat# P7170) prior to incubation with the following primary antibodies: HA (Biolegend, cat# 901501), PRRC2B (Santa Cruz Biotechnology, cat# sc-393604), tubulin (Sigma, cat# T9026T9026), vinculin (Sigma, cat# V9131), GAPDH (Merck, cat# MAB374), EIF3D (Bethyl, cat# A301-758A), EIF4G2 (BD Bioscience, cat# 610742), LARP1 (Proteintech, cat# 13708-1-AP), LARP4, (Bethyl, cat# A303-900A), PABP (Santa Cruz Biotechnology, cat# sc-32318), PRMT1 (Cell signaling, cat# 2449S), calnexin, (Abcam, cat# ab22595), RPS3, (Proteintech, cat# 66046-1-Ig), RPS14 (Abcam, cat# ab174661), and Biotin (Cell signaling, cat# 5597S). Secondary antibodies were either horseradish peroxidase (HRP)-conjugated goat anti-mouse (Jackson ImmunoResearch Labs, cat# 115-035-003) or anti-rabbit (Jackson ImmunoResearch, cat# 111-165-144), detected by enhanced chemiluminescence using SuperSignal™ West Pico PLUS Chemiluminescent Substrate (Thermo Scientific, 34580).

### Mass spectrometry

HA-PRRC2B_L_ and HA-PRRC2B_S_ and control HA-mCherry immuno-precipitates were subjected to in-solution tryptic digestion, or for methylation analysis, chymotrypsin digestion, and analyzed by LC/MS as previously described (*56*). Samples were loaded using split-less nano-Ultra Performance Liquid Chromatography (10 kpsi nanoAcquity; Waters, Milford, MA, USA) coupled online through a nanoESI emitter (10 μm tip; New Objective; Woburn, MA, USA) to a quadrupole orbitrap mass spectrometer (Q Exactive Plus, Thermo Scientific) using a FlexIon nanospray apparatus (Proxeon). Data was acquired in data dependent acquisition (DDA) mode, using a Top10 method. MS1 resolution was set to 70,000 (at 200m/z), mass range of 375-1500m/z, AGC of 1e6 and maximum injection time was set to 60 msec. MS2 resolution was set to 17,500, quadrupole isolation 1.7m/z, AGC of 1e5, NCE of 27%, dynamic exclusion of 25 sec and maximum injection time of 60 msec. Raw data was processed with MaxQuant v1.6.6.0 (*57*). The data was searched with the Andromeda search engine against the human (*Homo sapiens*) protein databases as downloaded from Uniprot (www.uniprot.com), appended with common lab protein contaminants as previously described (*56*). The quantitative comparisons were calculated using Perseus v1.6.0.7 (*58*). Decoy hits were filtered out and only proteins that were detected in at least two replicates of at least one experimental group were kept. Empty intensity values went through imputation using the corresponding Perseus function. A Student’s t-Test, after logarithmic transformation, was used to identify significant differences between the experimental groups, across the biological replica. Fold changes were calculated based on the ratio of geometric means of the different experimental groups. Interactions were considered significant for proteins with at least 2 peptides, whose abundance was enriched in PRRC2B_L_ or PRRC2B_L_ immune-precipitates over mCherry immune-precipitate more than 2-fold, FDR<0.05.

For post-translation modification analysis, raw data was processed with Proteome Discoverer v2.4 informatics platform (Thermo Scientific) and searched against the human protein database as above. The search was done with the SequestHT and Andromeda 2.0 search engines. Search parameters were defined to include chymotryptic peptides with up to two missed cleavages allowed. Fixed modification was set to carbamidomethylation of cysteines. Variable modifications were set to oxidation of methionines, protein N-terminal acetylation, and arginine methylation. Peptide and protein identifications were filtered at FDR of 1% using the Percolator software.

### Immunofluorescence staining

U2OS G3BP-GFP cells were plated in 6 well dishes for transfection with the indicated plasmids. 24 h post transfection the cells were trypsinyzed and replated on ibidi *μ*-Slide 8 Well plates (ibidi, cat# 80806). The following day the cells were treated with sodium arsenite (NaAsO_2_) (Sigma, cat# 71287) for 30 min. The cells were washed with PBS and fixed using 4% PFA (Electron Microscopy Science, 15710). Permeabilization was performed with PBS + 0.1% Triton X-100 (PBT), followed by blocking in PBS containing 5% Donor goat serum (NGS) (Biological Industries, cat# 04-009-1A) for 1h. Wells were incubated with primary antibody, mouse anti-HA diluted in blocking solution for 1 h at room temperature followed by 1 h with Alexa Fluor® 555 goat anti-mouse IgG, and then stained with DAPI. The wells were kept in Fluoromount G (Southern Biotech, cat# 0100-10) and visualized with Zeiss LSM900 confocal microscope, 40x C-Apochromat water-immersion objective, N.A. 1.2. Images were processed using ZEN software (version 2.4). For co-localization with mitochondria, HeLa cells grown on ibidi *μ*-Slide 8 Well plates were transfected with HA-PRRC2B_L_ and stained with MitoTracker™ Red CMXRos (Invitrogen, cat# M7512) for 30 minutes and HA as described above.

### RNA deep sequencing

RNA-seq libraries were prepared at the Crown Genomics institute of the Nancy and Stephen Grand Israel National Center for Personalized Medicine, Weizmann Institute of Science. Libraries were prepared using the TruSeq Stranded mRNA-seq kit (Illumina) according to the manufacturing protocol. Briefly, the polyA fraction (mRNA) was purified from 500 ng of total input RNA from PRRC2B KO and control NT cells grown in DMEM or EBSS media for 48h, followed by fragmentation and the generation of double-stranded cDNA. After Agencourt Ampure XP beads cleanup (Beckman Coulter), A base addition, adapter ligation and PCR amplification steps were performed. Libraries were quantified by Qubit (Thermo Fisher scientific) and TapeStation (Agilent). Sequencing was done on Novaseq Plus X 1.5B 300 cycles kit, allocating ∼50M reads per sample (Illumina; paired end sequencing).

Adapter trimming, alignment to hg38, and statistical analyses were performed with the UTAP pipeline (*59*), which utilizes STAR version 2.7.10a, cutadapt version 4.1 DESeq2(v1.36.0) for normalization and differential expression analysis. Importantly, batch was included as a co-factor in the statistical model. Additional DESeq2 parameters included: betaPrior=TRUE, cooksCutoff=FALSE, and independentFiltering=FALSE. The sva R package was used to apply batch correction to the log2-normalized counts, and the resulting output was used for PCA analysis. The data was analyzed first with all 4 samples, and then with only EBSS samples, as total RNA content was greatly reduced in these cells compared to DMSO. Criteria for passing the differential expression test were: BaseMean > 10, │log2FC│ > 0.263 (FC>1.2) and *p* < 0.05.

### Statistical analysis

Statistical analysis of large datasets (MS, RNA-seq) and sequence analysis is provided in the respective sections of the Materials and Methods above. Additional analysis as indicated in figure legends was done using GraphPad Prism 9 by paired two-tailed Student T-tests, with *p* < 0.05 considered statistically significant.

## Acknowledgements

We would like to thank Anastasia Lev for technical assistance with polysome purification and the staff of the G-INCPM, Weizmann Institute of Science, for preparing RNA libraries and performing RNA-seq experiments.

## Author Contributions

Conceptualization: NG, SB, AK

Data curation: AS, TO

Formal analysis: AS, TO, SP

Funding acquisition: AK

Investigation: NG, DB, SP, ME

Methodology: NG

Project administration: NG

Resources: AS, TO, ME

Supervision: AK

Validation – NG, SB

Visualization – NG, SB, ME, AS, TO, SP

Writing – original draft – NG, SB

Writing – review & editing – all authors

## Data and materials availability

The mass spectrometry proteomics data have been deposited to the ProteomeXchange Consortium via the PRIDE (*60*, *61*) partner repository with the dataset identifier PXD057527 and PXD057790. The RNA-seq datasets generated in the current study are available in the Gene Expression Omnibus (GEO), accession ID GSE282113. All other data are available in the main text or the supplementary materials.

