## Supplemental Figures S1-S3 for "The RNA-binding protein PRRC2B preserves 5’ TOP mRNA during starvation to maintain ribosome biogenesis during nutrient recovery"

Nadav Goldberg *et al.*

**This PDF file includes:**

Figs. S1 to S3

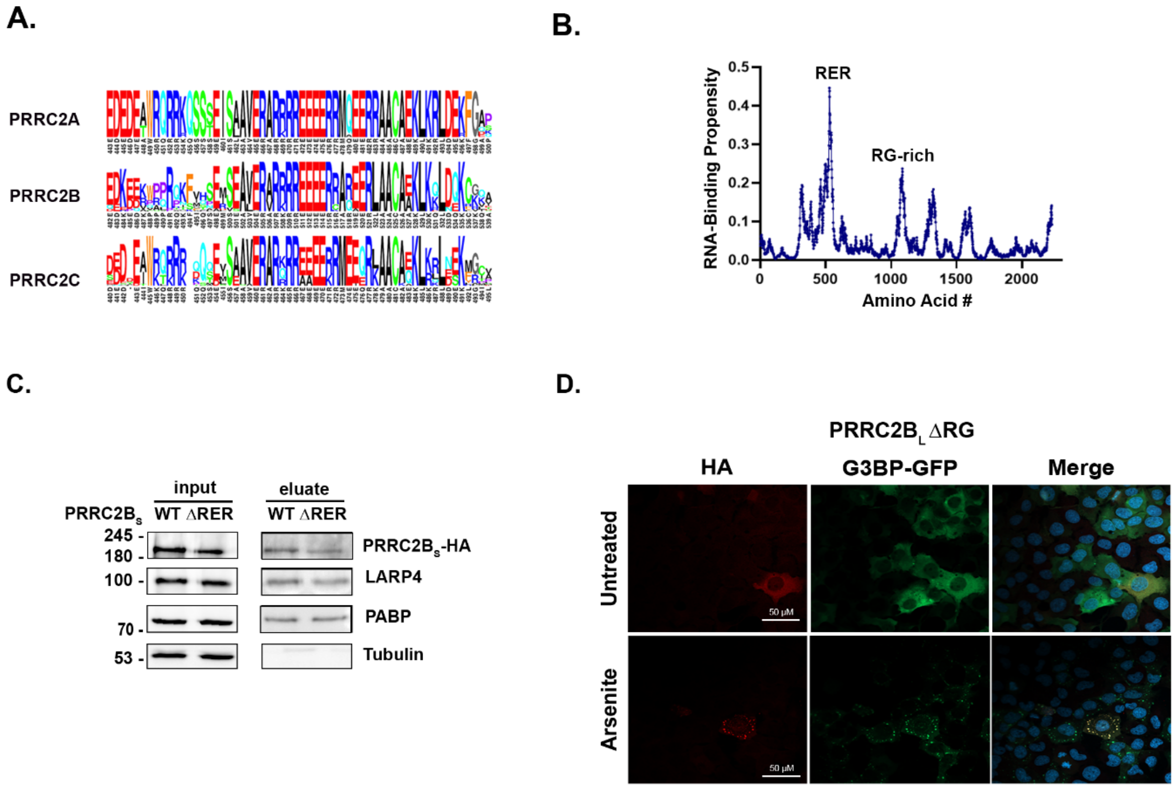


### Fig. S1.

**Additional analysis of PRRC2B isoforms.** (**A**) Multiple sequence alignment within the RER domain for all PRRC2B family members, PRRC2A, B, and C, shown according to the human amino acid position numbers. Significance was determined by Mann-Whitney U test for whether the sites outside exon 16 are less conserved than those in it, after weight-correction for occurrence of RG sites in human proteins; $p$=0.037. (**B**) Prediction of RNA binding capacity of PRRC2B according to DisoRDPbind. Each residue is assigned a value between 0 and 1 representing its likelihood to be involved in disordered RNA binding. Peaks correspond to regions likely to bind mRNA. Positions of RER and RG-rich domain are indicated. (**C**) Oligo-dT pull down assay on HEK 293T lysates expressing HA-tagged PRRC2B$_S$ WT or deleted of the RER domain. Lysates were cross-linked to stabilize RNA-protein complexes, and mRNAs with bound proteins were then captured on oligo-dT beads. Proteins present in the total cell lysate (input) and extensively washed bead eluate were probed with anti-HA antibodies to detect PRRC2B. Endogenous LARP4, PABP and Tubulin were used as positive and negative controls, respectively. (**D**) U2OS cells stably expressing GFP-G3BP were transfected with HA-tagged PRRC2B$_L$ deleted of the RG-rich domain and treated with sodium arsenite or DDW for 30 min. Cells were fixed and stained with anti-HA and DAPI to detect nuclei. (Related to Figures 1,2)

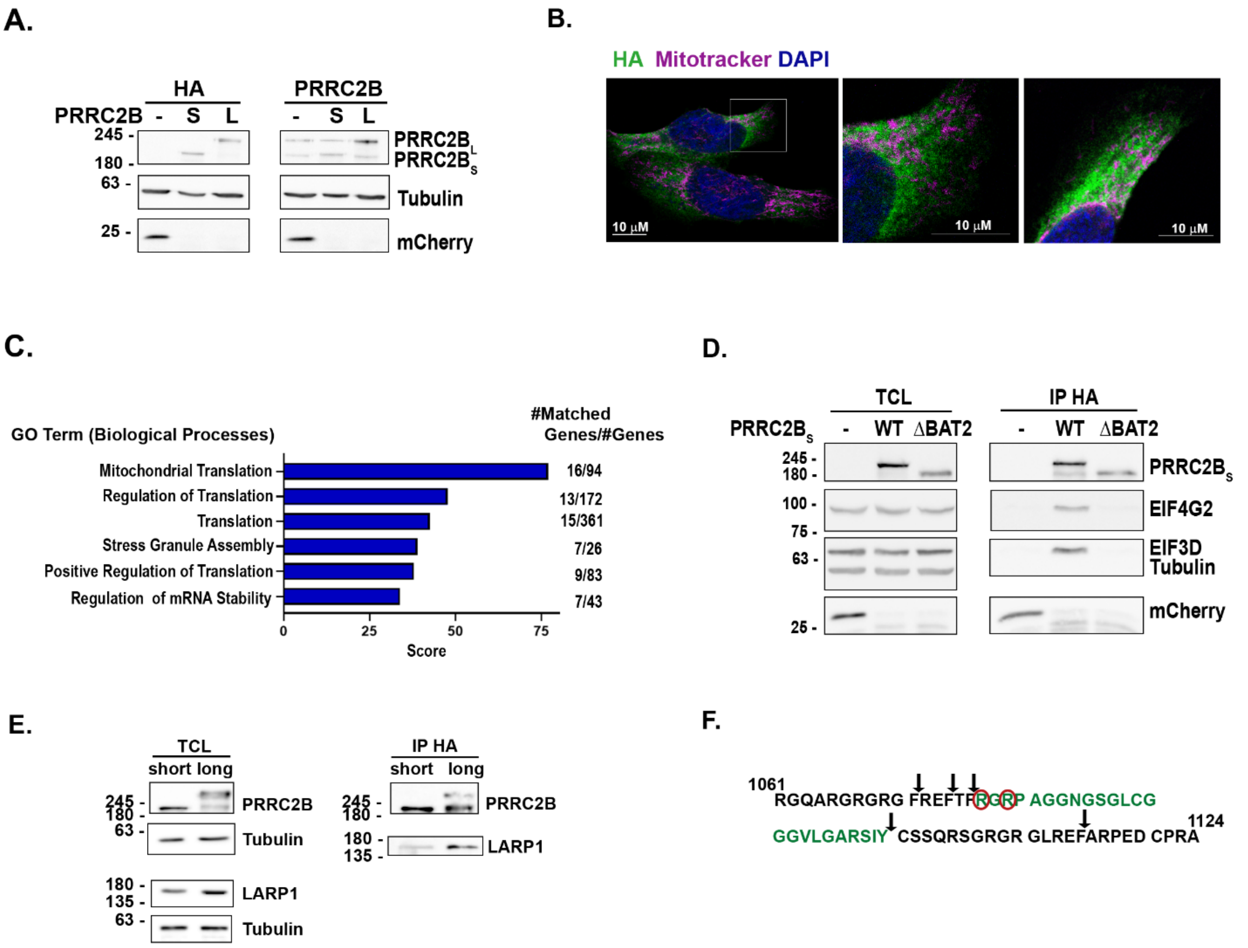


### Fig. S2.

**Additional analysis of PRRC2B$_L$ and PRRC2B$_S$ interactomes and localization.** (**A**) Representative western blot showing expression levels of exogenous HA-tagged mCherry, PRRC2B$_L$ and PRRC2B$_S$ in HEK 293T cells under same conditions as used for IP-Mass Spec experiment. (B) HeLa cells transiently transfected with pcDNA plasmid expressing HA-PRRC2B$_L$ were stained with anti-HA antibodies, Mitotracker to detect mitochondria and DAPI for nuclei. Middle panel is enlargement of boxed area in left panel. No overlap between HA and Mitotracker signals is observed. (**C**) Top 6 significant GO terms (Biological Processes) and scores identified by GeneAnalytics analysis of the set of 69 proteins that preferentially interact with PRRC2B$_L$. The number of matched genes within the set out of the total number of genes in the particular biological process is indicated at right of graph. (**D**) HEK 293T cells transiently transfected with pcDNA plasmids expressing HA-Cherry, HA-tagged PRRC2B$_S$ full length or ΔBAT2 mutant were immunoprecipitated with anti-HA antibodies, and lysates and IPs subjected to western blot analysis for the indicated proteins. (**E**) HEK 293T cells transiently transfected with pcDNA plasmids expressing HA-tagged PRRC2B$_L$ or PRRC2B$_S$ were immunoprecipitated with anti-HA antibodies, and lysates and IPs subjected to western blot analysis for the indicated proteins. Note that lower band in PRRC2B$_L$ lanes is a degradation product of the full length protein. (**F**) Sequence of RG-rich domain indicating Arg residues identified as methylated by Mass Spec following chymotrypsin digestion of the protein. Peptide identified by Mass Spec is colored green, remaining peptides were not detected. Chymotrypsin cleavage sites are indicated by arrows, methylated residues are circled in red. (Related to Figure 3).

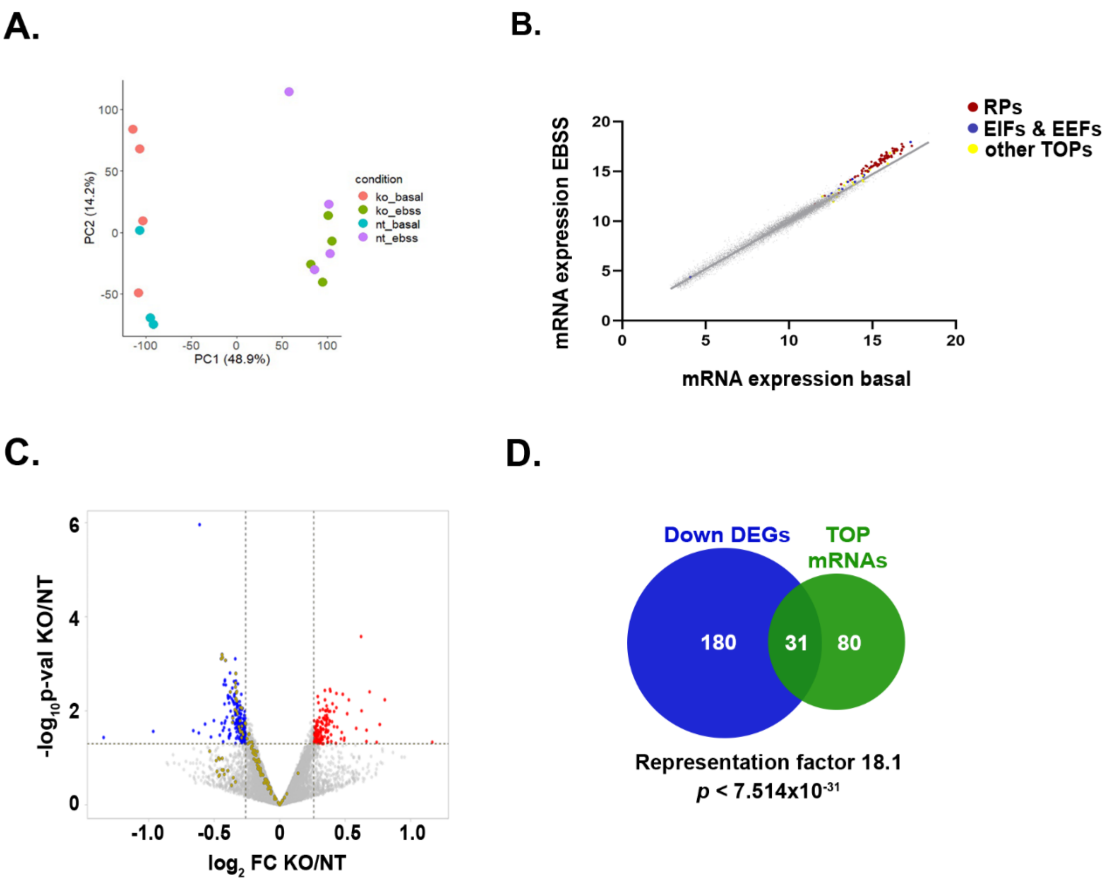


**Fig. S3.**

**Additional analysis of RNA-seq data in starved PRRC2B cells.** (**A**) Principle Component Analysis (PCA) of RNA seq data from NT or PRRC2B KO, basal or EBSS conditions. (**B**) Normalized expression levels of total mRNAs under basal conditions vs starvation (EBSS), for protein-coding genes only. mRNAs above the grey line are expressed at higher relative levels during starvation compared to the rest. TOP mRNAs are highlighted, specifically ribosomal proteins and eukaryotic initiation and elongation translation factors (EIFs, EEFs). (**C**) Volcano plot of the fold-change in mRNA expression levels in PRRC2B KO vs NT control under starvation conditions, vs. their significance expressed as $-\log_{10}$ *p*-value. Increased and decreased differentially expressed genes are indicated in red and blue, respectively. Yellow dots represent mRNAs encoding TOP mRNAs. (**D**) Venn diagram showing overlap between TOP mRNAs and the group of statistically significant down-regulated genes (down DEGs). (Related to Figure 4)
